# A reference genome for the Andean cavefish *Trichomycterus rosablanca* (Siluriformes, Trichomycteridae): building genomic resources to study evolution in cave environments

**DOI:** 10.1101/2023.11.11.566715

**Authors:** Carlos Daniel Cadena, Laura Pabón, Carlos DoNascimiento, Linelle Abueg, Tatiana Tiley, Brian O-Toole, Dominic Absolon, Ying Sims, Giulio Formenti, Olivier Fedrigo, Erich D. Jarvis, Mauricio Torres

## Abstract

Animals living in caves are of broad relevance to evolutionary biologists interested in understanding the mechanisms underpinning convergent evolution. In the Eastern Andes of Colombia, populations from at least two distinct clades of *Trichomycterus* catfishes (Siluriformes) independently colonized cave environments and converged in phenotype by losing their eyes and pigmentation. We are pursuing several research questions using genomics to understand the evolutionary forces and molecular mechanisms responsible for repeated morphological changes in this system. As a foundation for such studies, here we describe a diploid, chromosome-scale, long-read reference genome for *Trichomycterus rosablanca*, a blind, depigmented species endemic to the karstic system of the department of Santander. The nuclear genome comprises 1Gb in 27 chromosomes, with a 40.0x HiFi long-read genome coverage having a N50 scaffold of 40.4 Mb and N50 contig of 13.1 Mb, with 96.9% (Eukaryota) and 95.4% (Actinopterygii) universal single-copy orthologs (BUSCO). This assembly provides the first reference genome for the speciose genus *Trichomycterus*, which will serve as a key resource for research on the genomics of phenotypic evolution.

## Introduction

Upon colonizing cave environments, a variety of animals (various arthropods, crustaceans, several vertebrates) have converged in morphology, physiology, and behavior (Juan *et al*. 2010; Romero 2011; Protas and Jeffery 2012). While common selective pressures leading to adaptation presumably account for such evolutionary convergence, the loss of traits in cave organisms has also been attributed to genetic drift (Wilkens and Strecker 2017; Policarpo *et al*. 2021). The remarkable degree to which different traits have evolved convergently and in concert also suggests that various constraints may place limits on adaptation and drive the course of evolution in cave systems (Franz-Odendaal and Hall 2006). At the molecular and developmental level, phenotypic changes associated with living in caves could be accounted for either by mutations in coding regions of genes (Kim *et al*. 2011; Warren *et al*. 2021) or by changes in patterns of gene expression (van der Weele and Jeffery 2022; Arcila *et al*. 2023), possibly involving phenotypic plasticity (Bilandžija *et al*. 2020) and epigenetics (Gore *et al*. 2018).

In fish, species in multiple evolutionary lineages have evolved convergently in caves (Chakrabarty *et al*. 2012; Armbruster *et al*. 2016; Hashemzadeh Segherloo *et al*. 2018). Losses or reductions in eyes and pigmentation are especially well known (Niemiller *et al*. 2019), but whether similar phenotypes of different fish species living in caves have evolved using equivalent molecular mechanisms remains to be determined. Also, existing information is insufficient to establish the relative importance of changes in coding regions of genes versus changes in regulatory regions that change patterns of gene expression to account for phenotypic evolution. Developing new genomic resources is fundamental to pursue research in such systems.

We recently assessed whether fishes in the Neotropical genus *Trichomycterus* (Siluriformes) inhabiting cave environments in the Eastern Cordillera of the Andes of Colombia and which lack eyes and pigmentation evolved via a: (1) single event of colonization of subterranean environments and subsequent vicariance or dispersal leading to the origin of new species; or (2) multiple colonizations of caves from surface environments followed by evolutionary convergence (Flórez *et al*. 2021). Employing mitochondrial DNA sequences to infer phylogeographic relationships, we found that caves in this region have been colonized separately by at least two different clades of *Trichomycterus*. In addition, we documented shallow to non-existent mtDNA divergence between surface and cave populations, even though they differ considerably in morphology; this suggests that their divergence is recent or has proceeded in the face of gene flow, with selection counteracting homogenizing effects of migration (Flórez *et al*. 2021). In either case, the system is an attractive model to assess the genetic basis of adaptation to life in caves using genomic tools.

We report a high-quality *de novo* reference genome generated for the cave specialist species endemic to Colombia *Trichomycterus rosablanca* using long-read sequencing and the Vertebrate Genomes Project (VGP) genome assembly pipeline. This reference genome will serve as a vital resource for studies aimed at understanding the genetic basis of phenotypic evolution in cave environments.

## Methods

### Sampling tissue

We collected tissue samples from a male individual of *T. rosablanca* captured in June 2021 by MT, CDC and Maykol Galeano at the type locality of this recently described species (Mesa S. *et al*. 2018), namely La Sardina Cave in El Peñón, Santander, Eastern Cordillera of the Andes of Colombia (6º 05’ 36.0’’ N, 72º 49’ 42.7’’ W; Figure 1). Tissues were flash-frozen in liquid nitrogen in the field and stored at -80°C at Universidad de los Andes, prior to sending samples for laboratory work to the Rockefeller University Vertebrate Genome Lab. Our fieldwork was made possible by a collecting permit issued by the Colombian Ministry of Environment to the Universidad de los Andes (project PR.6.2020.7867, resolution 1177, October 9, 2014 - IDB 0359), and we exported samples following authorization by the Agencia Nacional de Licencias Ambientales (export permit 02529). The *T. rosablanca* voucher specimen was deposited in the fish collection at Instituto Alexander von Humboldt under catalogue number IAvH-P 28386.

**Figure 1.**
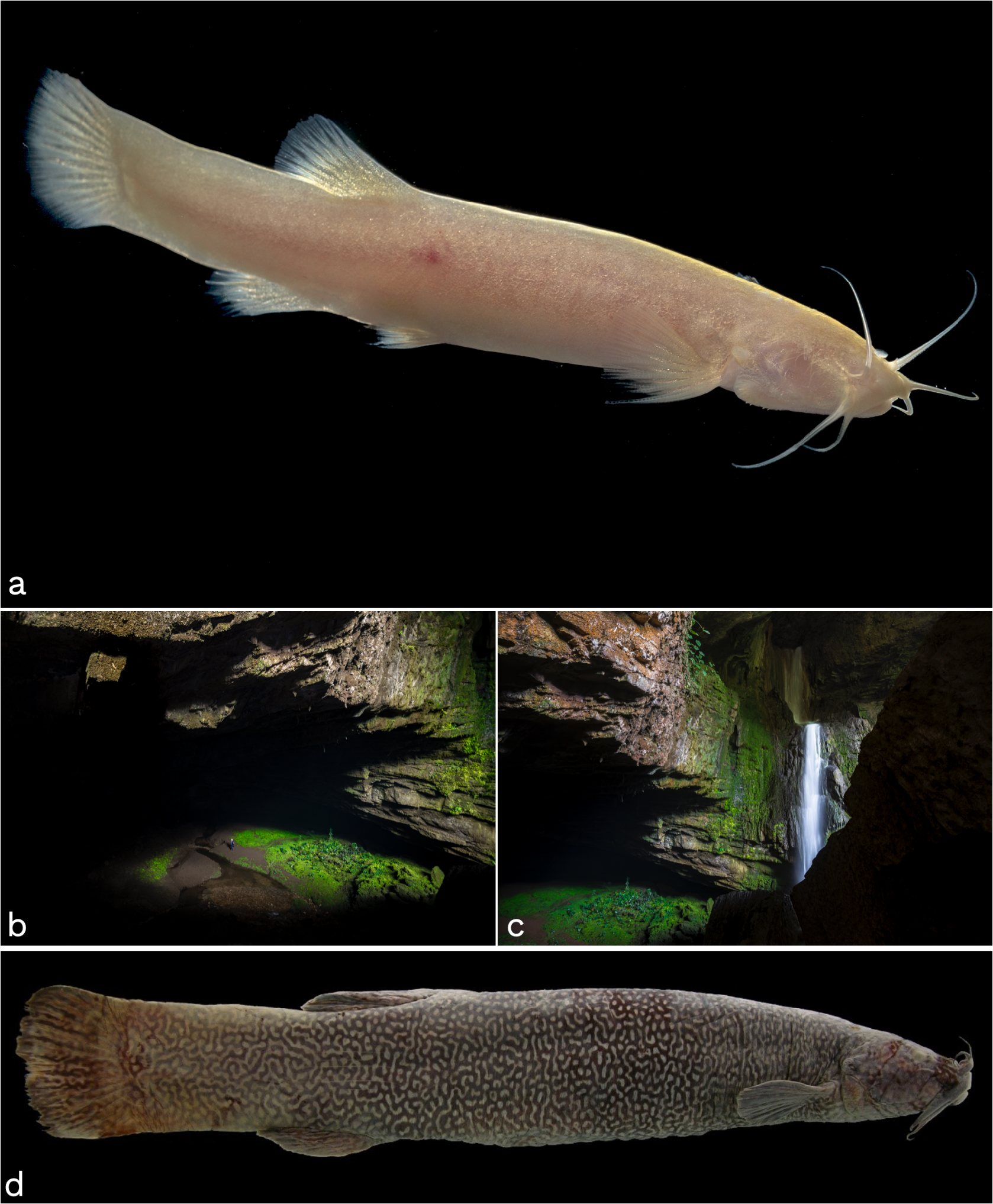
*Trichomycterus rosablanca* is a Neotropical catfish species lacking pigmentation and eyes, restricted to karstic cave environments in the Colombian Andes. (a) Specimen of *T. rosablanca* from El Peñón, Santander, Colombia. (b, c) Views of La Sardina Cave, the type locality of *T. rosablanca* where we conducted fieldwork for this study. (d) The putative sister species of *T. rosablanca* is *Eremophilus mutisi*, a heavily pigmented, sighted species living in surface rivers and streams in the Eastern Andes of Colombia. Photographs by Felipe Villegas (a-c) and Carlos DoNascimiento (d).

### DNA extraction and sequencing

For DNA extraction, an input of 50 mg of fin tissue was disrupted with the TissueRuptor II (Qiagen Cat. No.9002755) and high molecular weight DNA was isolated using the MagAttract HMW DNA Kit (Cat. No.67563). The DNA was not sheared prior to PacBio library preparation. We used 5 μg of purified DNA to prepare a PacBio smrtbell library with the SMRTbell prep kit 3.0 (Pacific Biosciences 102-141-700) and size-selected with the AmpurePB bead method to remove all templates smaller than 5,000 bp. The library was quantified with a Qubit 3 fluorometer (Invitrogen Q33216) and its size was assessed with the Agilent Femto Pulse. The library was sequenced on a Sequel IIe instrument with Sequencing Plate 2.0, Binding Kit 3.2, and two 8M SMRT cells, to generate a total of 40X coverage (20X per haplotype) of Pacbio high-fidelity (HiFi) long reads. To generate Hi-C data, we used the Arima-HiC 2.0 kit (Arima Genomics, Carlsbad, CA, USA) on a liver sample, according to the manufacturer’s protocols. This protocol generates Hi-C cros-links in the tissue before library preparation and sequencing. The library was sequenced on an Illumina NovaSeq 6000 platform with 2x150bp read length at Psomagen, Inc. (Rockville, MD, USA).

### Assembly

To assemble the chromosome-scale reference genome, we followed the VGP 2.0 diploid pipeline in Galaxy (Larivière *et al*. 2023) using the Rockefeller University instance on a high-performance computing system. Contigs were generated using HiFiasm (v0.16.1+galaxy4) in Hi-C phasing mode (PMID: 35332338). Both haplotypes (Hap1 and Hap2) were individually scaffolded using yahs (v1.2a.2+galaxy0) and the Hi-C data (Zhou *et al*. 2023). Both haplotypes underwent a decontamination protocol to remove any extraneous bacterial or viral sequences, using the NCBI decontamination pipeline. Hap1 then underwent manual curation to fix any noted structural errors and naming chromosomes. We also generated the complete 16,581 bp mitochondrial genome using MitoHiFi (PMID: 37464285). Summary statistics were generated with gfastats (Formenti *et al*. 2022).

### Transcriptome sequencing

To generate transcriptome data for annotation, RNA-Seq data was generated from the same animal used for the genome, but from brain, muscle, liver, and testes tissue. mRNA preparation was polyA selected, libraries prepared in paired-end layout using the Illumina Stranded mRNA Prep, and sequencing conducted on a Illumina NovaSeq 6000. Sequence output ranged from 65 to 168 million reads per sample. Raw data have been deposited in NCBI SRA, under the VGP Transcriptome Sequencing BioProject PRJNA516733 (SRA Accession #s SRX22284548 for brain, SRX22284547 for muscle, SRX22284540 for liver, SRX22284539 for testes.

### Analysis of BUSCO genes

We evaluated the completeness of expected gene content based on BUSCO (Manni *et al*. 2021) with the Eukaryota_odb10 and Actinopterygii_odb10 data sets.

### Data availability

All sequencing data and assembly are available from NCBI BioProject PRJNA924296. The whole-genome sequence is deposited in NCBI under accession numbers GCA_030014385.1 (Hap1) and GCA_030015355.1 (Hap2), and named fTriRos1.Hap1 and fTriRos1.Hap2, respectively. These assemblies and the mitochondrial genome are also in GenomeArk (https://www.genomeark.org/genomeark-all/Trichomycterus_rosablanca.html)

## Results and Discussion

HiFiasm in Hi-C mode on the Pacbio HiFi reads to generate haplotype-phased contigs, followed by yahs and Hi-C for scaffolding, and manual curation using Hi-C contact maps, led to highly contiguous and relatively complete assemblies for all chromosomes in both haplotypes (Supplementary Fig 1). Haplotype 1 was chosen as the reference assembly, as it was the most complete (Supplementary Table 1), with the final assembly size of 1.04 Gb, the longest contig length of 87 Mb, contig N50 of 13.1 Mb, and contig L50 of 24. Manual curation identified 27 chromosomes (Fig. 2a).

**Figure 2.**
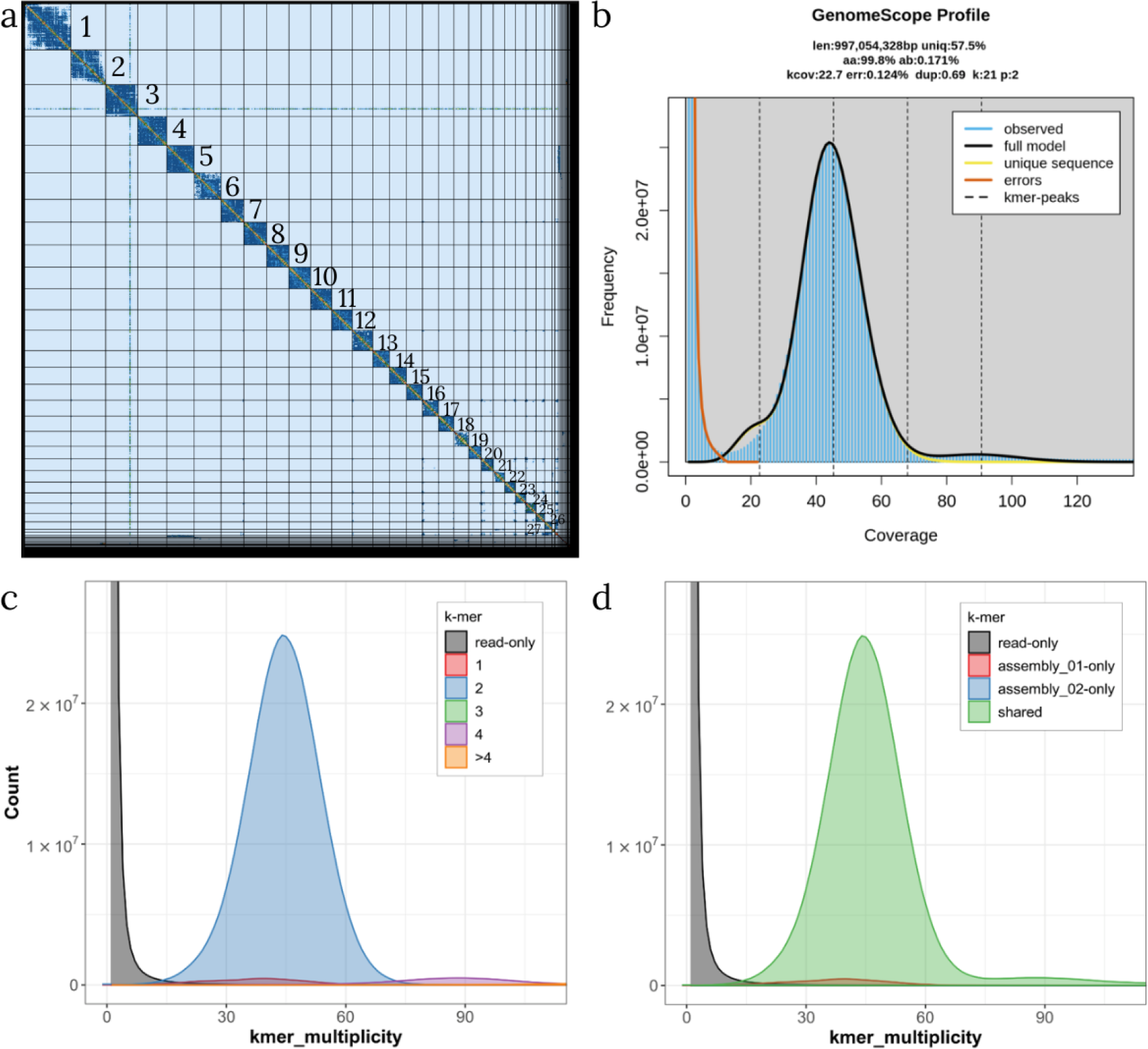
(a) Hi-C contact map of the Hap1 assembly of *Trichomycterus rosablanca* after manual curation with the corresponding chromosome number placed on the side of each portion of the map. (b) GenomeScope2 profiling summary of the diploid genome (p:2), with the estimated genome size (len), percentage of heterozygosity based on k-mer distributions. (c) Merqury copy numbers spectrum plot. (d) Merqury assembly spectrum plot for evaluating k-mer completeness.

Actinopterygii BUSCO gene representation was 96.7% complete with 95.4% single-copy genes, while the eukaryote BUSCO gene representation was 99.6% complete with 96.9% single-copy genes. The assembly has a GC content of 40.27% (Table 1). Details of the assembly statistics for both haplotypes are reported in Supplementary Table 1.

**Table 1.** Basic assembly statistics, BUSCO statistics run with actinopterygii_odb10 containing 3640 BUSCO groups and Merqury statistics for the *Trichomycterus rosablanca* genome.

|  |  |  |
| --- | --- | --- |
| **Assembly statistics** | Assembly length | 1,045,025,885 bp |
|  | Number of scaffolds | 349 |
|  | Scaffold N50 | 40,4 Mb |
|  | Scaffold L50 | 10 |
|  | Number of contigs | 547 |
|  | Contig N50 | 13.1Mb |
|  | Contig L50 | 24 |
| **BUSCO** | Complete (C) | 96.7% |
|  | Complete single copy (S) | 95.4% |
|  | Complete duplicated (D) | 1.3% |
|  | Fragmented (F) | 1.1% |
|  | Missing (M) | 2.2% |
| **Merquy** | Completeness | 99.40% |
|  | QV | 58.8169 |

There was 0.171% of nucleotide heterozygosity between the two haplotypes, as predicted in GenomeScope k-mer distribution analysis, with a genome size estimation of 997,054,325 bp and 0.124% error rate (Fig 2b). Merqury calculated an overall assembly completeness of haplotype 1 of 99.40% and QV score of 58.8169, with a higher percentage of completeness in haplotype 1 compared to haplotype 2 (S Table 2). Merqury copy numbers spectrum and assembly spectrum plots show that most of the reads are in both haplotypes, while most of the unique reads are found in Hap1 (Fig 2c-d).

Our research program plans to use genomic resources developed for *Trichomycterus* cavefish and their surface counterparts to understand genetic and developmental mechanisms involved in the repeated origin of troglomorphic phenotypes in this system. We first plan to analyze the genome of *Trichomycterus* species with typical cave and surface phenotypes in a comparative framework with genomic information available for fishes in other lineages and other organisms which have evolved similar phenotypes independently after experimenting similar selective pressures (Kim *et al*. 2011; McGaugh *et al*. 2014; Manni *et al*. 2019; Warren *et al*. 2021). We expect such comparisons will allow us to elucidate whether there are mutations in coding regions potentially responsible for phenotypic adaptation in *Trichomycterus* or if explanations based on changes in regulatory regions and patterns of gene expression should be pursued. In case mutations in regions associated with phenotypic variation (i.e. genes involved in eye formation or in melanin deposition in the skin) are detected, comparisons with other organisms will allow us to ask whether convergent phenotypic evolution in different branches of the tree of life has employed similar or different mechanisms (Elmer and Meyer 2011; Bo *et al*. 2022).

We are also working on a population-scale study based on reduced-representation approaches, whole-genome resequencing and transcriptomics hoping to achieve a better understanding of fine-scale phylogeography and spatial patterns of variation in neutral and functionally important genetic diversity in Colombian *Trichomycterus*. This will be coupled with in-depth studies of population-level variation in phenotypic traits describing body shape and sensory organs not previously studied in surface and cave *Trichomycterus*. As with similar work in emerging model systems like *Astyanax* cavefish (Moran *et al*. 2023), we expect our ongoing analyses to shed light on the genetic underpinnings of phenotypic variation and on the role of processes of dispersal and various forms of selection in driving phenotypic evolution.

## Acknowledgements

We thank Maykol Galeano for help with fieldwork and Mireya Osorio for assistance with permits. This project was funded by the Facultad de Ciencias at Universidad de los Andes through a *Programa de Investigación* awarded to CDC (Convocatoria 2021-2023). EDJ’s effort was supported by the Howard Hughes Medical Institute.

---

**Supplementary Figure 1.**
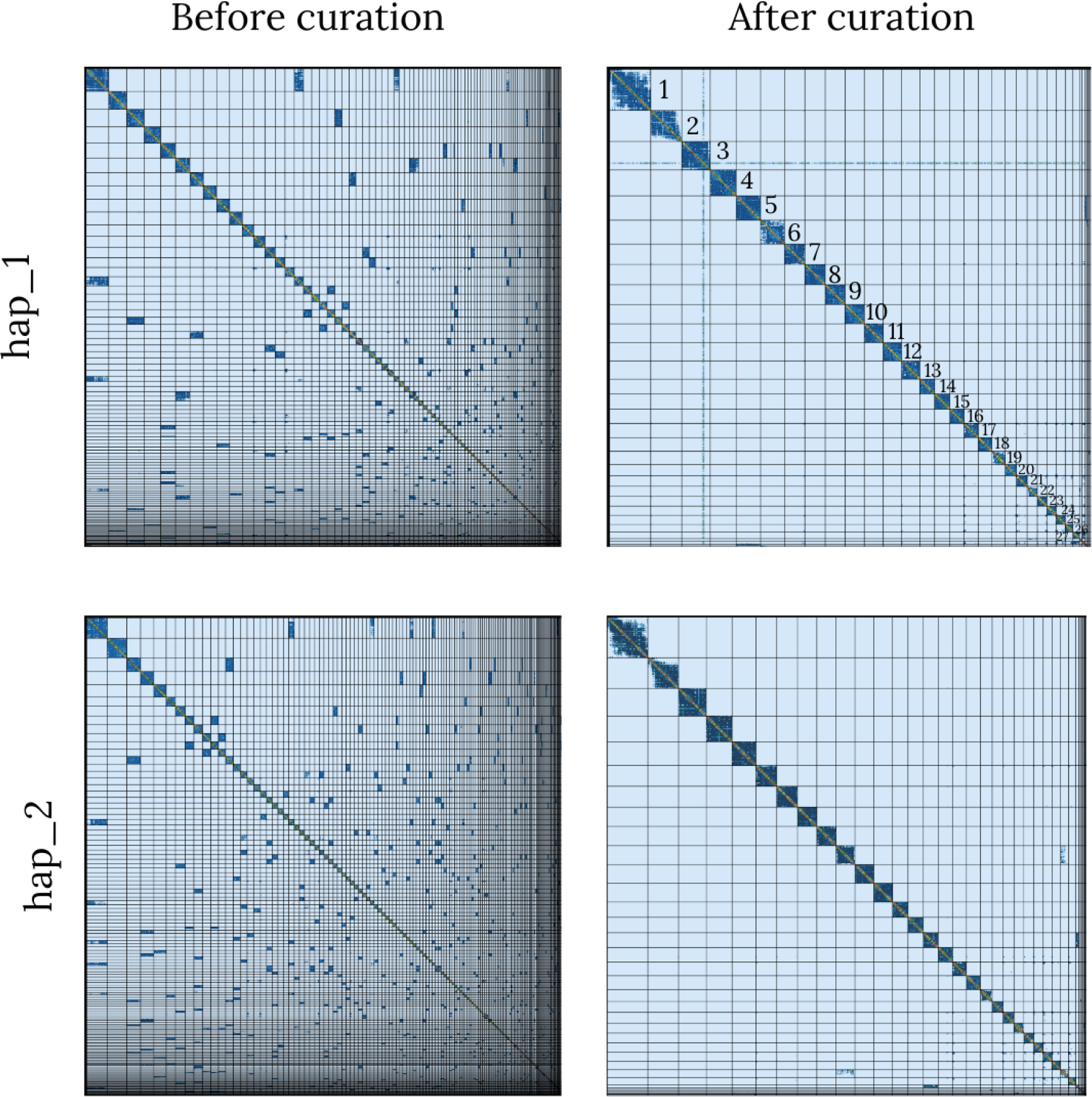
HiC contact maps for hap_1 and hap_2 before and after curation. Hap_1 was manually curated leading to a chromosome-level assembly.

**Supplementary Table 1.** Assembly statistics for *Trichomycterus rosablanca* assembly of Hap_1 and Hap_2. BUSCO statistics run against 3354 BUSCO groups of the Actinopterygii lineage.

|  |  | **Hap_1** | **Hap_2** |
| --- | --- | --- | --- |
| **Assembly statistics** | Assembly length | 1,045,025,885 bp | 994462882 |
|  | Number of scaffolds | 417 | 297 |
|  | Average Scaffold length | 2511521.94 | 3348359.87 |
|  | Scaffold N50 | 40182557 | 3893692 |
|  | Scaffold L50 | 10 | 10 |
|  | Scaffold NG50 | 40182557 | 38085262 |
|  | Scaffold LG50 | 10 | 11 |
|  | Largest scaffold | 85,843,722 | 82356993 |
|  | Smallest scaffold | 1000 | 1000 |
|  | Number of contigs | 551 | 489 |
|  | Average contig length | 1900685.75 | 2033587.9 |
|  | Contig N50 | 13074164 | 8272383 |
|  | Contig L50 | 24 | 34 |
|  | Contig NG50 | 13266506 | 8272383 |
|  | Contig LG50 | 22 | 34 |
|  | Largest contig | 40,042,000 | 39880030 |
|  | Smallest contig | 1000 | 1000 |
| **BUSCO** | Complete (C) | 3273 | 3161 |
|  | Complete single copy (S) | 3235 | 35 |
|  | Complete duplicated (D) | 38 | 22 |
|  | Fragmented (F) | 23 | 136 |
|  | Missing (M) | 58 | 3354 |

**Supplementary Table 2.** Merqury assembly statistics for *Trichomycterus rosablanca* assembly of Hap_1 and Hap_2.

|  | **Hap_1** | **Hap_2** | **Both** |
| --- | --- | --- | --- |
| **Solid k-mers in assembly** | 621116666 | 60814703 | 621855155 |
| **Solid k-mers in reads** | 625587997 | 625587997 | 625587997 |
| **Completeness %** | 99.29 | 97.21 | 99.40 |
| **Unique k-mers** | 28629 | 27671 | 56300 |
| **Common k-mers** | 1047268109 | 994415582 | 2041683691 |
| **QV** | 58.8547 | 58.7776 | 58.8169 |
| **Error rate** | 1.30177e-06 | 1.32508e-06 | 1.31313e-06 |

